# Cell-specific isolation and functional analysis of primary cardiomyocyte-derived large extracellular vesicles from infarcted mouse hearts

**DOI:** 10.64898/2026.10.01.755891

**Authors:** Lélia Borowski, Alina Ramazanova, Paul Alayrac, Vincent Duval, Morgane Aubert, Léa Pineau, Louis Faranton, Camille Knosp, Estelle Robidel, Florent Dingli, Damarys Loew, Jose Vilar, Chantal M. Boulanger, Xavier Loyer, Jean-Sébastien Silvestre, Stéphane M. Camus

## Abstract

Extracellular vesicles (EVs) are membrane-bound particles secreted by all cells and serve as critical mediators of intercellular communication in both physiological and pathological contexts. However, elucidating the precise functions of endogenous EVs remains challenged by a fundamental technical bottleneck: current isolation methodologies depend on shared physical properties or generic surface markers, preventing the selective enrichment of cell-type-specific EVs within complex tissues. Consequently, EVs recovered from biofluids or tissue extracts represent heterogeneous populations, confounding the direct attribution of biochemical signatures and biological activities to distinct cellular origins.

Here, we describe a versatile genetic-labeling and affinity-based purification strategy that enables the selective isolation of cell-specific primary EV. Using cardiomyocyte-derived EV (CM-EVs) as a proof-of-concept, we crossed Myh6-Cre mice with the *ROSA26-*mTmG reporter line to restrict membrane-targeted GFP expression to cardiomyocytes. Following myocardial infarction, cardiac EV were recovered and GFP-positive CM-EVs were selectively purified by anti-GFP nanobody-based magnetic pulldown. This approach yields highly enriched, functional cardiomyocyte-derived extracellular vesicles (CM-EVs). To accommodate diverse analytical needs, we established two distinct elution workflows: (i) a denaturing protocol optimized for downstream proteomics and immunoblotting, and (ii) a gentle, non-denaturing protocol that preserves vesicle integrity and biological activity for downstream *in vitro* and *in vivo* functional assays.

Using this method, we characterized CM-EVs size distribution and proteomic signature, and demonstrated their capacity to transfer functional cargo to target cells *in vitro*. Because the purification approach relies solely on targeting a genetically restricted membrane tag rather than assumed surface markers, it is immediately adaptable to virtually any cell type for which a validated Cre driver exists. This method removes a long-standing bottleneck in EV research and provides a broadly applicable platform for deciphering the biochemical composition and in-vivo functions of cell-type-specific primary EV in physiological and pathological settings.

## Introduction

Extracellular vesicles (EVs) are nanoscale, membrane-bound particles secreted by virtually all cell types. Once dismissed as mere cellular waste, EVs are now established as crucial drivers of intercellular communication across physiological processes, such as development and immune regulation, as well as pathological states, including cancer and atherosclerosis (1) (2) (3). By carrying diverse molecular cargo, including proteins, lipids, and nucleic acids, EVs dynamically reflect the identity and functional status of their parent cells (4). Within the cardiovascular system, EVs derived from cardiomyocytes (5, 6), endothelial cells, fibroblasts, and immune cells orchestrate myocardial homeostasis, inflammatory cascades, and tissue remodeling (3, 7). Furthermore, their enhanced stability in circulation relative to labile signaling molecules (e.g., cytokines) highlights their potential as robust biomarkers for cardiovascular disease.

Yet, despite this promise, deciphering the precise functional roles of EV has been slowed by a persistent technical limitation: EV are highly heterogeneous, and current isolation methods cannot resolve or purify EV populations according to their cell of origin. EV are commonly categorized into small EV (multivesicular bodies-derived exosomes, <200 nm) and large EV (plasma membrane-derived vesicles, 200–1000 nm), but widely used separation methods such as differential ultracentrifugation, density gradients, size-exclusion chromatography or immunoprecipitations yield mixtures of vesicles with overlapping sizes, densities, and surface features. Consequently, EV isolated from tissues or biofluids represent composite populations originating from numerous cell types. This complexity is particularly pronounced in organs such as the heart, where cardiomyocytes, fibroblasts, endothelial cells and infiltrated immune cells all release EV during homeostasis and in response to injury.

Attempts to selectively capture specific EV subsets through immunoaffinity enrichment have offered only limited progress. Most established EV markers, including the tetraspanins CD9, CD63 and CD81, are shared across multiple EV subtypes and cell types, precluding cell-type specificity (8, 9). Moreover, immunocapture often traps EV irreversibly or requires harsh, detergent-based elution that disrupts vesicle structure and prevents downstream functional assays. Even recent innovations in elution-compatible immunocapture remain constrained by their reliance on universal markers_(10). Fundamentally, no endogenous surface antigen is known to uniquely identify EV from a single cell type within complex tissues. As a result, assigning cell-type-specific functions to primary EV populations *in vivo* has remained largely inaccessible.

To overcome this technical bottleneck, we established a proof-of-concept genetic labeling and affinity-purification platform that enables the selective isolation of cell-type–specific EVs directly from intact tissue microenvironments. As a model system, we crossed Myh6-Cre driver mice with ROSA26-mTmG reporter mice to restrict membrane-targeted GFP expression specifically to cardiomyocytes. Because mGFP incorporates into cardiomyocyte plasma membranes, it is subsequently embedded into the limiting membrane of cardiomyocyte-derived EVs (CM-EVs). Consequently, GFP-tagged CM-EVs within cardiac tissue lysates can be selectively enriched using high-affinity anti-GFP nanobodies conjugated to magnetic beads.

We further developed two elution workflows that expand the versatility of this method: (i) a denaturing elution optimized for biochemical analyses such as label-free proteomics and immunoblotting; and (ii) a gentle, non-denaturing elution that preserves vesicle architecture, enabling downstream *in vitro* and *in vivo* functional assays. Together, these steps constitute a robust pipeline for isolating, enriching, and interrogating primary CM-EVs in physiological or pathological contexts.

Here, we present and thoroughly validate this GFP-based isolation strategy, demonstrating that it yields highly specific, structurally intact CM-EVs capable of functional cargo delivery. Crucially, this platform is universally adaptable: by pairing any validated Cre-driver line with mTmG reporters, researchers across disciplines can now isolate and interrogate cell-type–specific EVs from virtually any lineage within complex tissue environments. By overcoming a decades-long technical bottleneck, this method provides the field with a powerful framework to decipher the precise molecular architectures and biological functions of endogenous EVs in homeostasis, inter-organ crosstalk, and disease progression.

## Materials and Methods

### Transgenic mice

We used Myh6-Cre mice (B6.Cg-Tg(Myh6-cre)2182Mds/J) crossed with the ROSA26-mTmG reporter line (B6.129(Cg)-*Gt(ROSA)26Sor*^*tm4(ACTB-tdTomato,-EGFP)Luo*/J). In the double-transgenic offspring (Myh6-Cre mTmG), Cre recombination in cardiomyocytes excises the floxed tdTomato cassette and activates GFP expression. Membrane targeting of the GFP protein is achieved by fusing it to the modified N-terminal domain of the plasma membrane-anchored protein Myristoylated Alanine-Rich C-kinase Substrate (MARCKS). All experiments were conducted according to the ethical committee for animal experimentation (University of Paris-Cité, CEEA 34) and the National Charter on the ethics of animal experimentation from the French Ministry of Higher Education and Research under the reference MESR # D75-15-32.

### Myocardial infarction model

Myocardial infarction was induced in adult Myh6-Cre mTmG mice (10–14 weeks old) via permanent ligation of the left anterior descending (LAD) coronary artery. Mice were anesthetized with an intraperitoneal injection of ketamine (100 mg/kg; Ketamine 1000, Virbac) and xylazine (10 mg/kg; Rompun 2%, Bayer), and maintained on a homeothermic blanket system (Harvard Apparatus) throughout the procedure. A left thoracotomy was performed through the fourth intercostal space, the pericardium was carefully removed, and the LAD was ligated with a 7-0 monofilament suture (Peters Surgical). Following surgery, mice were allowed to recover on a 37°C heating pad after extubation. Experimental groups were matched for age and littermates and mice were randomly enrolled to either MI surgery group or control group.

### Two-photon microscopy

Mice were injected with a lethal dose of pentobarbital sodium and perfused with ice-cold PBS. Hearts, spleens and lungs were harvested, washed in ice-cold PBS and fixed in 4% paraformaldehyde (PFA) for 30 min at 4°C, protected from light. Samples were then washed in PBS and stored at 4°C until processing. Pieces of approximately 1 mm³ were cut from each organ with a sterile scalpel. For endogenous fluorescence imaging, tissues were permeabilized in PBS containing 0.5% Triton X-100 for 1 h at room temperature. After three washes in PBS, tissues were incubated PBS supplemented with Hoechst 33342 (20 µg/mL, B2261, Sigma-Aldrich) for 30 min at room temperature. After three PBS washes, tissues were mounted on microscope slides fitted with silicone isolators (GBL665104, Sigma-Aldrich) within Mowiol 4-88 mounting medium (81381, Sigma-Aldrich) and imaged on a Leica SP8 DIVE multiphoton microscope with spectral detection (4Tune module) through a 25× water-immersion objective (Fluotar VISIR, NA 0.95). Hoechst, membrane-targeted GFP (mGFP) and membrane-targeted tdTomato (mTomato) were excited sequentially at 700, 920 and 1040 nm, respectively. Multichannel images (1024 × 1024 pixels) were saved as TIFF files in LAS X software (Leica).

### Isolation of primary cardiomyocytes

Adult mouse cardiomyocytes were isolated by enzymatic perfusion using a Langendorff-free method, as previously described (11). All solutions were sterile-filtered through a 0.2 µm membrane and prewarmed to 37°C before use. Mice were anesthetized with a lethal dose of pentobarbital sodium and intravenously injected with 20 µL of heparin prior to exsanguination. Hearts were perfused *in situ* with PBS through the right ventricle. The ascending aorta was clamped, and hearts were harvested and transferred into Petri dishes containing ice-cold sterile PBS. Hearts were then cannulated through the left ventricle and sequentially perfused with warm digestion buffer containing collagenase type II and type IV (Worthington) and protease type XIV (Sigma-Aldrich), delivered continuously at 80 mL/h for 30 minutes. Atria and right ventricles were removed and ventricles were minced with a sterile scalpel. Enzymatic activity was stopped by addition of FBS-based stop solution and the cell suspension was filtered through a 100 µm cell strainer and gently homogenized. Cardiac cell suspension was allowed to sediment for 15 min at room temperature and plated in 6 wells culture plates. Cellular fluorescence was subsequently assessed immediately after plating using a 20X objective (0.45 NA, Fluorite, LWD) on an EVOS FL fluorescence microscope (Thermo Fisher Scientific) fitted with GFP (AMEP 4651, Invitrogen) and Texas Red (AMEP 4655, Invitrogen) filter cubes.

### Isolation of EV from infarcted heart tissue

EV were isolated from cardiac left ventricles according to a previously described procedure (6). Briefly, hearts were perfused with ice-cold PBS and harvested in sterile saline (0.1 µm-filtered 0.9% NaCl) to isolate the left ventricles. The left ventricles were then minced with sterile scissors for 30 seconds on ice. To remove cell debris, the samples were vortexed and centrifuged twice at 400 × g for 15 minutes at 4°C. The resulting EV-containing supernatants were centrifuged at 20,500 × g for 45 minutes at 4°C. The pellets, enriched in large extracellular vesicles (lEV), were resuspended in 200 µL of sterile saline (0.1 µm-filtered 0.9% NaCl), aliquoted and stored at −80°C until use.

### H9C2 CD9-eGFP cell culture

Rat cardiomyoblasts H9C2 stably overexpressing human CD9 inserted in the plasmid **pCL6-** CD9eGFP (H9C2-CD9-eGFP) (generous gift from Dr A. Görgens) were cultured in DMEM (30-2002, ATCC) supplemented with 10% fetal calf serum and 1% penicillin/streptomycin. Cells were maintained at 37°C in a humidified atmosphere containing 5% CO₂ and cultured in non-coated flasks. Cells were maintained until approximately 80% confluence and then passed. The culture medium was renewed every 2 days, and cells between passages 4 and 7 were used for experiments.

### Isolation of EV from H9C2 CD9-eGFP cultured medium

H9C2 EVs were isolated from 48h conditioned media by differential centrifugation. The first centrifugation (600g, 15 min at 4°C) removed floating cells and debris while the second (4,500g, 20 min, 4°C) removed apoptotic bodies. The resulting supernatant was then transferred into ultracentrifugation tubes (3138-0050, Thermo Scientific) and centrifuged at 20,500g for 2 h at 4°C (Sorval Super T21, ST21, Kendro Laboratory) to pellet lEVs. lEVs were resuspended in 200 µL of sterile saline solution (0.9% NaCl filtered through a 0.1 µm membrane), aliquoted, and stored at −80°C until further use.

### GFP-trap immunocapture and CM-EVs elution

Large EV were incubated with GFP-Trap magnetic beads (M-270, Proteintech), which use a camelid anti-GFP nanobody. Typically, 25 µL of bead suspension was washed with sterile saline (0.1 µm-filtered 0.9% NaCl) then added to the EV samples. The mixture was then incubated overnight at 4°C under agitation. Unbound EV were removed and beads were washed three times with sterile saline (0.1 µm-filtered 0.9% NaCl). Bound GFP+ CM-EVs were then eluted under different conditions, depending on the subsequent analysis. For western blot analysis, we used 30 µL of Laemmli buffer (161-0747, Bio-Rad) with 200 mM dithiothreitol (DTT, 11583786001, Sigma-Aldrich); elution was carried out at 95°C for 5 minutes. For mass spectrometry, bound CM-EVs were eluted using either 0.15% trifluoroacetic acid (TFA; 28904, Thermo Fisher Scientific) or a Triton X-100-based buffer (150 mM NaCl (Sigma-Aldrich), 50 mM Trizma base, pH=8 (Sigma-Aldrich) and 1% Triton X-100 (Sigma-Aldrich)) before denaturation at 95°C for 10 minutes. For functional experiments, CM-EVs were either eluted in 1 M Trizma base (Sigma-Aldrich) at pH=2 or 12 for acidic or alkaline elutions, respectively, or in Glycine (Chromotek) or high ionic strength (5M LiCl2, pH=7, Sigma-Aldrich) elution buffers for 30 minutes at 37°C with constant agitation. Basic or acidic eluates were subsequently neutralized with the addition of concentrated hydrochloric acid or sodium hydroxide. Immediately after elutions, eluates were then placed on ice and stored at −80°C.

### Cardiac EV size distribution

EV size distribution was determined by tunable resistive pulse sensing (TRPS; Exoid, iZON) using an NP400 nanopore membrane suitable for particles ranging from 185 to 1100 nm (iZON). Size calibration was performed with CPC400 (mode diameter, 350 nm) calibration beads. EV concentrations were expressed as particles per milliliter.

### Flow cytometry determination of CM-EVs concentration

EV were analyzed by high-resolution flow cytometry (CytoFLEX, Beckman Coulter). Samples were diluted in sterile saline (0.9% NaCl filtered at 0.1 µm) on a per-sample basis to obtain an acquisition rate of ∼1000 events/s. The large-EV gate was set using calibration beads (Megamix-SSC, 0.16–0.5 µm; Biocytex). To determine purified CM-EVs concentration, samples were labeled with an anti-GFP antibody (chicken polyclonal, A10262, Thermo) for 30 minutes on ice. After ultracentrifugation (46,000 rpm for 1h at 4°C (Optima MAX-XP fitted with an MLA-130 rotor, Beckman Coulter), lEV were incubated with an anti-chicken secondary antibody conjugated to Alexa Fluor 488 (A32931, Thermo) for 30 minutes on ice. On some occasions, unlabeled bulk cardiac EVs or labeled CM-EVs were treated with 0.1% Triton X-100 or 0.5% SDS (Sigma-Aldrich).

### Western blotting

Purified CM-EVs, cardiac EV and organ lysates were prepared in Laemmli buffer (161-0747, Bio-Rad) supplemented with 200 mM DTT (11583786001, Sigma-Aldrich) and denatured at 95°C for 5 minutes. Lysates were subjected to SDS-PAGE and proteins transferred onto a nitrocellulose membrane (0.2 µm, 1620112, Bio-Rad). Membranes were incubated with primary antibodies (1-5 µg/mL), washed, and labeled with species-specific HRP-conjugated secondary antibodies (Jackson ImmunoResearch). Primary antibodies included: anti-GFP rabbit polyclonal (ab290, Abcam), anti-MYH6 mouse monoclonal (GT565, Invitrogen), anti-CD9 mouse monoclonal (ab395032, Becton Dickinson), and anti-Cre recombinase rabbit polyclonal (Poly908, BioLegend). Membranes were incubated with Enhanced Chemi-Luminescence (Clarity Western ECL, 170-5061, Bio-Rad) and images were acquired using a digital CCD imager (ImageQuant 800 Fluor, Cytiva). Densitometric quantification was performed with Fiji (NIH).

### Proteomics and mass spectrometry

Sample preparation. Protein precipitation was performed by incubating 20 µL of sample overnight at -20°C with 180 µL of 80% methanol, 0.1 M ammonium acetate (buffer 1). Precipitates were sedimented by centrifugation at 14,000 × g for 15 minutes at 4°C, and the supernatant removed. Precipitates were rinsed with 100 µL of buffer 1, and residual buffer removed by evaporation on a SpeedVac. Proteins were reduced with 10 µL of 5 mM DTT at 57°C for 1 h and alkylated with 2 µL of 55 mM iodoacetamide for 30 minutes at room temperature in the dark. Trypsin/Lys-C (Promega) was added at a 1:100 (wt:wt) enzyme:substrate ratio, 37°C overnight. Digested samples were loaded onto homemade C18 StageTips (three AttractSPE disks, #SPE-Disks-Bio-C18-100.47.20, Affinisep) for desalting. Peptides were eluted with 40/60 MeCN/H2O + 0.1% formic acid, vacuum-concentrated to dryness, and reconstituted in 10 µL of 0.3% (v/v) TFA before LC-MS/MS analysis.

### LC-MS/MS analysis

Online chromatography was performed with an RSLCnano system (Ultimate 3000, Thermo Scientific) coupled to an Orbitrap Exploris 480 mass spectrometer (Thermo Scientific). Peptides were trapped on a C18 pre-column (75 µm i.d. × 2 cm; nanoViper Acclaim PepMap 100, Thermo Scientific) with buffer A (2/98 MeCN/H2O, 0.1% (v/v) formic acid) at 2.5 µL/min over 4 minutes. Separation used a 50 cm × 75 µm C18 column (nanoViper Acclaim PepMap RSLC, 2 µm, 100 Å, Thermo Scientific) at 50°C with a linear gradient of 2% to 30% buffer B (100% MeCN, 0.1% formic acid) at 300 nL/min over 91 minutes. Full MS scans were acquired in the Orbitrap over m/z 375–1500 at a resolution of 120,000 at m/z 200. The 20 most intense ions were isolated and fragmented by higher-energy collisional dissociation (HCD) and acquired at a resolution of 15,000 with the AGC target set to 100%. Ions of charge state 2+ to 6+ were selected; normalized collision energy was 30% and dynamic exclusion 40 s.

### Isolation of primary peritoneal macrophages

Peritoneal macrophages (PEM) from mTmG mice were harvested by peritoneal washes. Briefly, mice were injected with a lethal dose of pentobarbital sodium (Euthasol, Virbac) before injection of 10 mL of PBS/BSA2% in the peritoneal cavity. After 5 minutes, the PBS/BSA2% was retrieved and centrifuged (500 x g, 10 min). The pellet was resuspended in sterile RPMI-1640 supplemented with 10% fetal bovine serum and 1% penicillin-streptomycin. The cell suspension was seeded in 8-well imaging slides (154453, Thermo) for 24h.

### Incubation of quantification of membrane-GFP⁺ macrophages

Cells (PEM) were incubated with purified CM-EVs at a ratio of 1:100 (macrophage:CM-EVs) for 24h. At the end of treatment, the cells were washed with warm PBS and cells were incubated in fresh growth media for 72h. Cells were then washed with ice-cold PBS and fixed with ice-cold 2% paraformaldehyde (047377.9M, Thermo) for 30 minutes at room temperature. Cells were then permeabilized and non-specific antibody interaction sites blocked (PBS, 0.5% Saponin, 2% BSA) and incubated with anti-GFP chicken polyclonal primary antibody (A10262, Thermo) overnight at 4°C. Cells were then washed (five times, PBS, 4°C), incubated with species-specific fluorophore-conjugated secondary antibodies (Thermo Fisher Scientific), washed again, mounted with ProLong Antifade Diamond (Thermo Fisher Scientific). Single-plane confocal images were acquired on a Leica SP8 with a 40× objective at 1024 × 1024 pixels (0.2405 µm/pixel), using identical acquisition settings across conditions. Hoechst, mGFP, and mTomato were acquired in separate channels. One field of view was analysed per animal (untreated, n = 5; vehicle, n = 10; EV-treated, n = 15; 30 fields, 875 cells). Images were analyzed using a custom Python pipeline (Python 3.11; scikit-image 0.26, SciPy 1.17, tifffile). Nuclei were segmented from Hoechst and cell-associated regions defined independently of mGFP expression using the Hoechst and mTomato channels together with mGFP; detailed segmentation procedures are described in Supplementary Methods. Background-corrected mGFP fluorescence was quantified as the 90th percentile of pixel intensities within the membrane compartment surrounding each nucleus. mGFP positivity was defined using a single threshold corresponding to the 99th percentile of the pooled control-cell distribution (untreated + vehicle; 543 cells), which was applied unchanged to all images. No treated-condition mGFP signal was used for segmentation or threshold determination. For statistics, the percentage of mGFP⁺ cells per field was treated as the experimental unit, with one field analyzed per animal. Groups were compared using Kruskal–Wallis analysis followed by two-sided Mann–Whitney U tests; Cliff’s delta was used as the effect size. Values are presented as mean ± SEM. Statistical analyses were performed using SciPy 1.17 and statsmodels 0.15.

### Statistical analysis

Data are presented as mean ± SEM. Statistical analyses were performed with Graphpad Prism software. Statistical tests are described in the figure legends. Sample sizes are described in the figure legends. P<0.05 was considered as significance threshold.

## Results

### Cardiomyocytes from Myh6-Cre mTmG mice specifically express GFP and generate GFP-positive large extracellular vesicles after myocardial infarction

To achieve cell-type–specific membrane tagging, we crossed Myh6-Cre driver mice with ROSA26-mTmG reporter mice. In these animals, Cre recombinase activity is restricted to cardiomyocytes, driving exclusive expression of membrane-targeted GFP (mGFP; Fig. 1A). Immunoblot analysis of tissue lysates confirmed that Cre and GFP were detected uniquely in Myh6-Cre mTmG cardiac tissues, but were absent in wild-type (WT) hearts and spleen, across both genotypes (Fig. 1B). Two-photon imaging of intact myocardium (Fig. 1C) and fluorescence microscopy of isolated primary adult cardiomyocytes (Fig. 1D) corroborated the strict, cardiomyocyte-specific localization of GFP.

**Figure 1.**
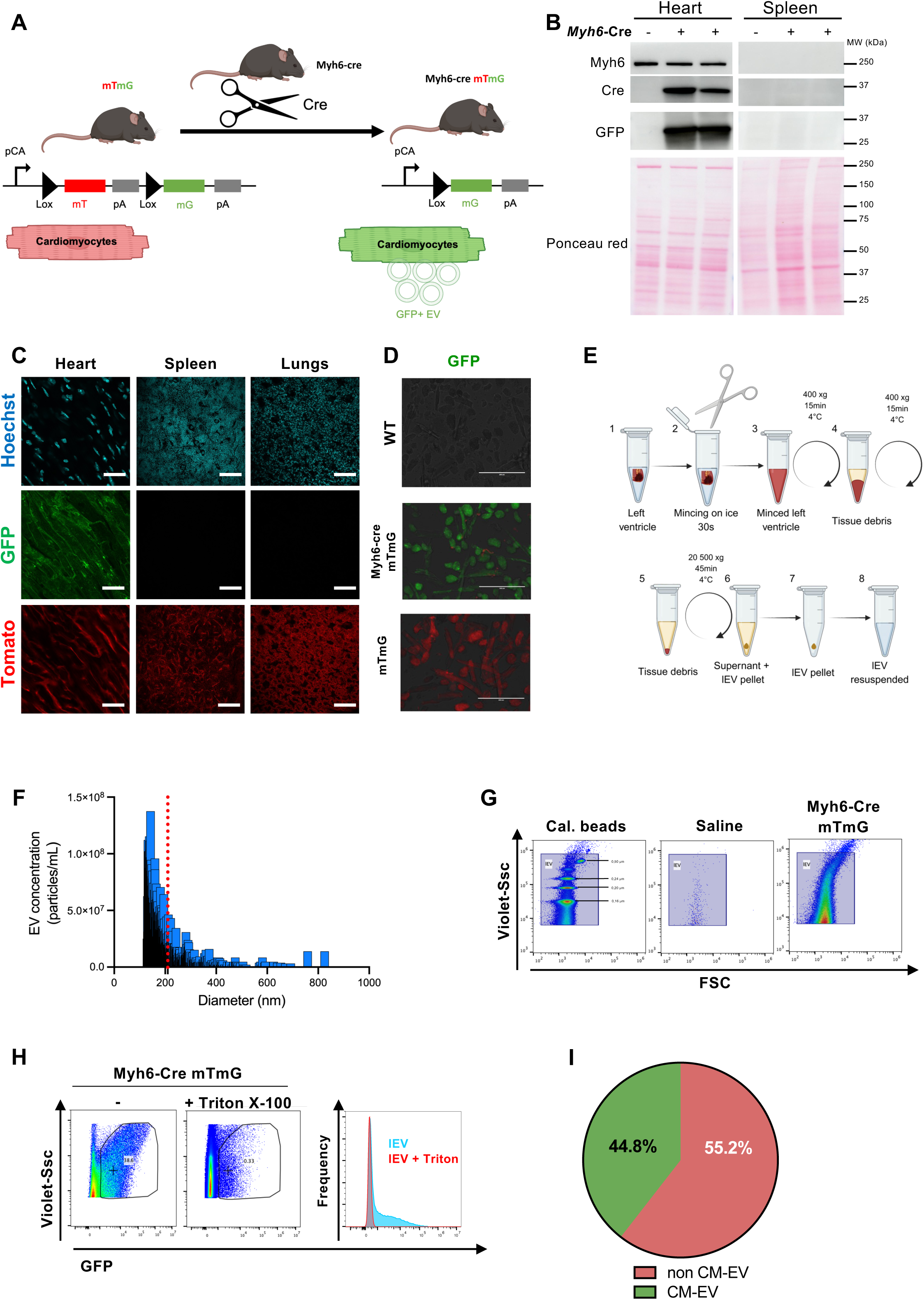
Cardiomyocytes from Myh6-Cre mTmG mice specifically express GFP and generate GFP-positive large extracellular vesicles after myocardial infarction. **A** Generation of Myh6-Cre mTmG transgenic mice by crossing Myh6-Cre driver mice with the Rosa26-mTmG lineage-tracing reporter line. **B** Cardiac and spleen lysates from Myh6-Cre mTmG (Cre+) and WT (Cre−) mice immunoblotted for MYH6, CRE recombinase and GFP. Ponceau red staining shows comparable protein loading. **C** Two-photon imaging of cardiac, spleen and lung slices from Myh6-Cre mTmG mice showing GFP expression (green) only in cardiomyocytes; tdTomato fluorescence (red); DAPI nuclear counterstain (blue). Scale bars: 20 µm. **D** Immunofluorescence of primary adult cardiomyocytes isolated from WT, Myh6-Cre mTmG and mTmG mice showing GFP-associated fluorescence only in Myh6-Cre mTmG cardiomyocytes. Scale bars: 200 µm. **E** Schematic of the cardiac large extracellular vesicle (lEV) isolation protocol. **F** Size distribution of isolated cardiac lEV measured by TRPS (Exoid) from Myh6-Cre mTmG hearts. The red dashed line indicates the mean particle diameter. (n = 3). **G** Flow cytometry of isolated cardiac lEV; calibration beads (0.16–0.5 µm) define the lEV gate, in which GFP+ lEV from Myh6-Cre mTmG hearts are detected. **H** Near-complete disappearance of GFP+ lEV events after 0.1% Triton X-100 treatment, confirming their lipid-vesicular nature. **I** Proportion of GFP+ cardiomyocyte-derived lEV among total cardiac lEV determined by flow cytometry (n = 3).

To evaluate whether cardiomyocyte-derived EVs (CM-EVs) retain GFP fluorescence, we subjected Myh6-Cre mTmG mice to experimental myocardial infarction (MI) via permanent ligation of the left anterior descending (LAD) coronary artery. Cardiac EVs were isolated from heart tissue 24 hours post-MI using differential centrifugation (Fig. 1E). Tunable Resistive Pulse Sensing (TRPS) of the isolated large EV (lEV) fraction revealed a particle size distribution spanning 100 to 700 nm, with a mean diameter of 213 nm (Fig. 1F), consistent with prior characterizations of cardiac lEVs (5). Flow cytometry analysis of the lEV pellet demonstrated distinct GFP-positive events within the gated lEV population (Fig. 1G). Detergent treatment attenuated these fluorescent events, confirming their lipid-bilayer nature (Fig. 1H). Quantitatively, GFP+ particles comprised an average of ∼45% of total isolated cardiac lEVs per heart (Fig. 1I). Together, these results confirm that Cre-mediated recombination is strictly restricted to cardiomyocytes and that GFP+ cardiomyocytes actively release fluorescently labeled lEVs following ischemic injury, validating this reporter system for selective EV capture.

### Purification of primary large cardiomyocyte-derived extracellular vesicles for biochemical characterization

Immunoblot analysis of cardiac lEV from Myh6-Cre mTmG and WT mice showed co-expression of the cardiomyocyte marker MYH6 and the EV marker CD9. GFP and CRE proteins were detected only in Myh6-Cre mTmG samples (Fig. 2A), confirming the presence of CM-EVs within the cardiac EV fraction. To specifically purify these CM-EVs, we subjected cardiac lEV preparations to a GFP-Trap magnetic pulldown (Fig. 2B). Immunoblot analysis of the eluates showed a 5-fold and a 2-fold increase in GFP and CD9 signal in the eluate, relative to input (Fig. 2C-E), validating the efficiency and specificity of GFP+ lEV purification.

**Figure 2.**
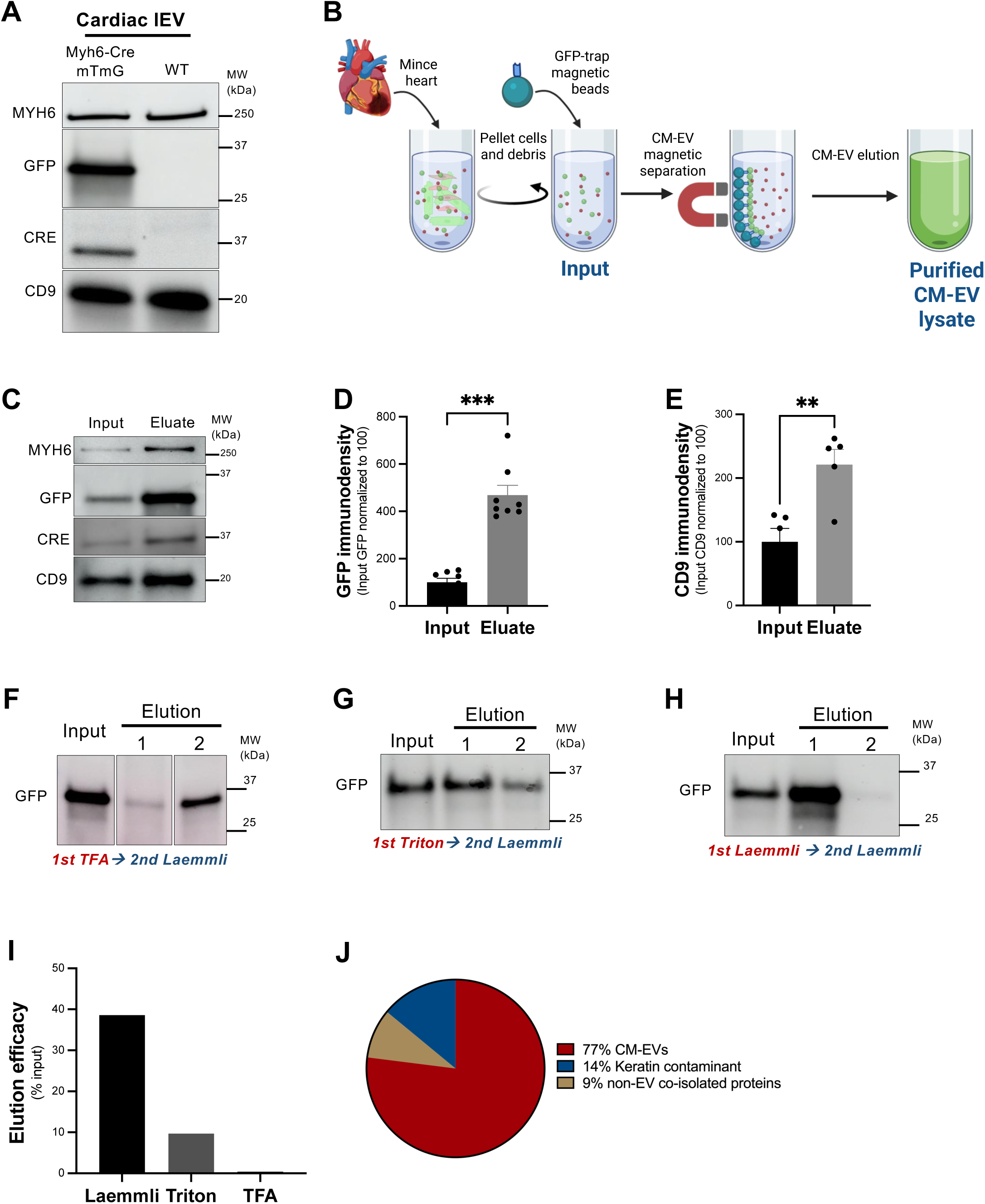
Purification of primary large cardiomyocyte-derived extracellular vesicles for biochemical characterization. **A** Immunoblot of cardiac lEV lysates from Myh6-Cre mTmG and WT mice; both express MYH6 and CD9, whereas the CRE recombinase and the GFP are detected only in Myh6-Cre mTmG lEV. **B** Schematic of the CM-EVs GFP-Trap purification workflow. **C** Representative immunoblots of cardiac input and purified CM-EVs (eluate) from Myh6-Cre mTmG hearts. **D** Purification yields a 5-fold increase in GFP (n = 8) and **E** a 2-fold increase in CD9 (n=5). **F-H** Representative immunoblots of purified CM-EVs eluted with TFA-based, Triton X-100-based or Laemmli-based buffers, each followed by a second Laemmli elution to assess residual bead-bound material. **I** Quantification of lEV protein elution efficacy calculated as immunodensity ratio between the elution and the input. **J** Proportions of purified CM-EVs proteins identified by mass spectrometry and classified by category (EV-associated, cardiomyocyte-specific, other cell source, non-vesicular).

To determine whether our purified CM-EVs preparations could be analyzed by proteomics, we adapted the elution step for downstream mass spectrometry (MS) and compared two elution buffers on their ability to elute the GFP: an acid-based buffer (trifluoroacetic acid, TFA) and a detergent-based buffer (Triton X-100 based buffer) (Fig. 2F and G, lanes 1).

To evaluate the efficiency of these elution buffers, we quantified the residual, un-eluted protein by subjecting the bead fraction to a second denaturing elution step (Laemmli sample buffer at 95 °C) (Fig. 2F-G, lane 2). These results were benchmarked against a control sample processed through two consecutive rounds of direct Laemmli elution (Fig. 2H, lanes 1 and 2).

The Triton buffer showed superior elution efficiency over the TFA buffer (Fig. 2F-I) and was therefore chosen for purified CM-EVs sample preparation. Label-free mass spectrometry of eluted CM-EVs identified 1,700 distinct proteins, the majority of which (∼77%) belonged to cardiomyocyte-derived (e.g., MYH6) and EV-derived (spanning the various categories from 1a to 5b as defined per the MISEV 2023) families. Contamination was minimal, with only 14% attributed to keratin contamination and 9% of the protein content to non-vesicular co-isolated material (Fig. 2J), underscoring the specificity and purity of the isolation. Together, these results demonstrate that our GFP+ lEV strategy provides specific, robust and reproducible isolation of primary CM-EVs from Myh6-Cre mTmG hearts.

### Purifying primary large cardiomyocyte-derived extracellular vesicles for functional characterization

To probe the pathophysiological role of CM-EVs, we first needed a way to recover them intact and at a usable concentration. We initially adapted a fluorescence-activated cell-sorting (FACS) strategy, gating on GFP to separate CM-EVs from the remaining cardiac EV (Fig. 3A, “purification strategy 1”). Sorting did recover GFP+ CM-EVs; however, because these vesicles are scarce relative to the bulk EV population, they were collected in a large volume and at a concentration far below that of the input, leaving the sorted material too dilute for any functional assay. We then attempted to perform ultracentrifugation in order to concentrate FACS-sorted CM-EVs, but this caused the loss of most of the sorted vesicles (Fig. 3B, C). The standard elution methods used for biochemical characterization are detergent-based and, by design, disrupt EV integrity to collect EV membranes, associated proteins, and luminal content. Releasing nanobody-bound EVs while maintaining their structural and functional integrity is notoriously challenging due to the exceptionally high affinity of the anti-GFP nanobody for its epitope. To overcome this limitation, we evaluated a panel of non-denaturing elution buffers spanning a range of ionic strengths and pH values (Fig. 3A, “Purification strategy 2”) for their ability to dissociate GFP from the magnetic beads. Starting with the 2 subsequent rounds of denaturating Laemmli elutions, we show that only one round of elution is enough to elute most of the CM-EVs attached to the magnetic beads, setting this type of elution as a benchmark for the other buffers (Fig. 3D). Next, the comparison of the different non-denaturating elution buffer showed that only the basic (high pH value) elution buffer showed some potency in eluting the GFP (Fig. 3E-I), defining our non-denaturating buffer of choice for subsequent purified CM-EVs functional testing. We also showed that the basic elution buffer was efficient enough for us to also detect both the GFP and the Cre recombinase in the eluate (Fig. 3J-L). Tunable Resistive Pulse Sensing (TRPS) comparison analysis between purified CM-EVs fraction and total cardiac lEV indicated a mean diameter size shift from 292 for purified CM-EVs, to 321 nm for total cardiac lEV, (Fig. 3M), not only confirming the physical integrity of CM-EVs but also their nature, as large EVs. Using this non-denaturating elution strategy, we then evaluated the integrity of eluted GFP+ CM-EVs by flow cytometry. We detected a marked increase in GFP+ events within the EV gate of the eluted CM-EVs sample compared to the input sample, indicating that the Basic elution buffer did not alter the GFP positivity and the morphological characteristics of eluted CM-EVs (Fig. 3N). Furthermore, we determined that the purification efficacy of this method was comparable between naïve or infarcted heart EV preparations (Fig. 3O). To confirm the lipid-vesicular nature of the isolated CM-EVs, we treated the eluted preparations with SDS (12). Detergent solubilization reduced the number of detectable GFP+ events by approximately 80%, supporting the vesicular nature of the captured particles, while a residual population of detergent-resistant events remained detectable (Fig. 3P, Q). Finally, to test whether these isolated CM-EVs remain biologically functional, we incubated CM-EVs purified from infarcted Myh6-Cre mTmG hearts with primary macrophages isolated from mTmG mice at a ratio of 100 CM-EVs per 1 macrophage (Fig. 3R). As purified CM-EVs expressed Cre recombinase (Fig. 3J and L), we reasoned that incubating them with mTmG macrophages could result in a transfer of the Cre recombinase protein and trigger gene recombination and GFP expression in the target cells. We stained macrophages for GFP and nuclei (Hoechst 33342). Following incubation with CM-EVs, we detected that around 20% of macrophages exhibited robust GFP signal, as compared to vehicle-treated or naive controls where none were detected (Fig. 3S, T). These findings deliver a critical proof-of-concept demonstrating that primary, cell-type–specific EVs can be isolated to near-homogeneity while retaining functional bioactivity for cellular uptake and signaling assays. Collectively, these data establish that our affinity-enrichment platform selectively isolates and concentrates intact CM-EVs while fully preserving their structural integrity and recipient cell delivery capacity.

**Figure 3.**
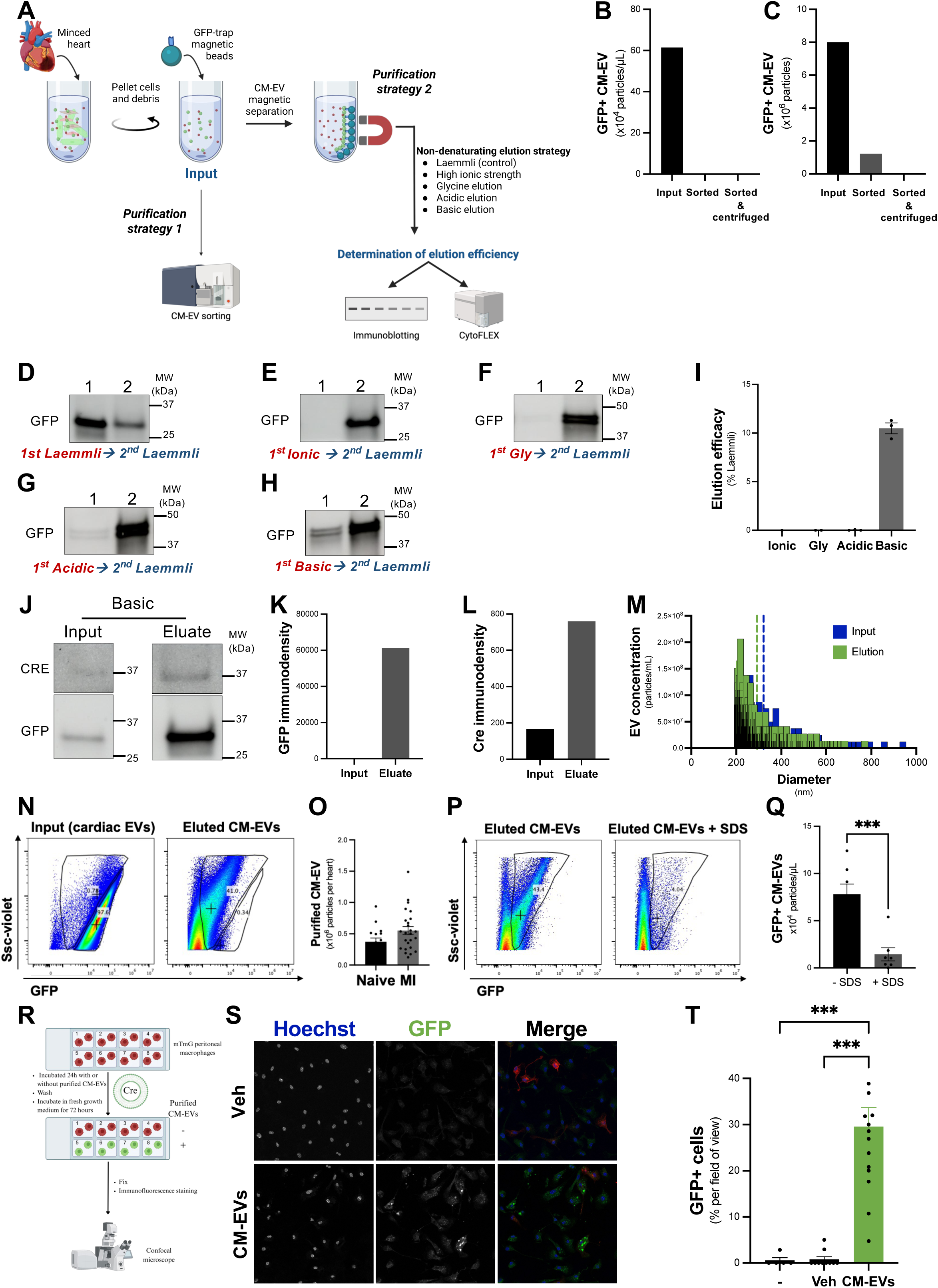
Purifying primary large cardiomyocyte-derived extracellular vesicles for functional characterization. **A** Schematic of the CM-EVs purification strategies tested. Quantification of FACS-sorted CM-EVs, expressed as GFP+ CM-EVs per µL (**B**) and as total GFP+ CM-EVs (**C**) before (input) and after sorting (sorted) and after concentration (sorted, centrifuged) (n = 1). **D–H** Representative immunoblots of purified CM-EVs eluted with different non-denaturing buffers (basic, acidic, glycine-based, and high ionic strength), compared with denaturing Laemmli buffer. **D–E** CM-EVs isolated from Myh6-Cre mTmG hearts; **F–H** EVs derived from H9C2 cells. For each condition, a second elution with Laemmli buffer was performed to assess residual bead-bound material. **I** The GFP protein was immunoblotted and immunodensity was quantified. (n = 3) **J** CM-EVs were eluted using the basic elution buffer, and the CRE recombisnase and the GFP were blotted. Immunodensities for the GFP (**K**) and the CRE recombinase (**L**) were quantified (n = 1). **M** Size distribution of isolated cardiac lEV (input, blue) and purified cardiomyocyte-derived lEV (Elution, green) isolated from Myh6-cre mTmG hearts and measured by TRPS (Exoid). The blue and green dashed lines indicate the mean particle diameter of cardiac lEV and purified cardiomyocyte-derived lEV populations, respectively (n = 1). **N** Flow cytometry assessment of purified CM-EVs after gentle elution with the basic elution buffer. Enriched, purified CM-EVs appear within the GFP+ EV gate. **O** Quantification of the average number of purified CM-EVs extracted per naïve (non-ischemic) or infarcted heart (n = 16 to 25). **P** Flow cytometry assessment of purified CM-EVs sensitivity to SDS. **Q** Quantification of purified CM-EVs before and after SDS incubation. (n = 7). **R** Schematic of macrophage incubation with purified CM-EVs to assess CM-EV-mediated transfer of active Cre recombinase. **S** Representative immunofluorescence images of macrophages incubated with purified CM-EVs from Myh6-Cre mTmG mice (Cre+) at a 100:1 (EV:macrophage) ratio or vehicle (CM-EVs suspension buffer). GFP and Hoechst nuclear staining. **T** Proportion of GFP+ mTmG peritoneal macrophages among total macrophages, per field of view. Four mice were used to isolate cells. Cells were seeded into 30 wells. n = 5, 10 and 15 wells per group, respectively. Mean ± SEM; Mann–Whitney U test. **p<0.01; ***p < 0.001.

## Discussion

Deciphering the pathophysiological contributions of primary, cell-type–specific EVs *in vivo* has long been constrained by a fundamental methodological bottleneck. While bulk EV isolation

strategies yield heterogeneous mixtures that mask cellular origin, existing functional studies rely almost exclusively on EVs harvested from cell cultures, an artificial setup that fails to recapitulate the complex signals of the native tissue niche. In this study, we overcome this longstanding limitation by establishing a genetically driven, affinity-based platform that selectively isolates EVs originating from a defined cell population directly from intact, diseased tissue. By capturing endogenous CM-EVs *in situ* under ischemic conditions, our approach bridges the gap between single-cell origin and tissue-level EV signaling, providing a scalable model to interrogate endogenous EV biology without the confounding artifacts of *in vitro* culture.

To address this limitation, we developed a primary EV isolation method based on genetic, cell-specific surface-tag expression followed by affinity capture, enabling the specific isolation of large EV (lEV) derived from cardiomyocytes *in vivo* during defined pathophysiological states. The method leverages the Myh6-Cre mTmG model, in which membrane-targeted GFP is selectively expressed in cardiomyocytes. By applying GFP immunocapture to heart-derived EV preparations from infarcted mice, we achieve robust isolation of a highly enriched cardiomyocyte-derived lEV population with minimal contamination. To our knowledge, this represents the first method enabling isolation of primary lEV from a single cell type *in situ* and *in vivo*, in a relevant pathophysiological setting.

A major limitation of EV research to date has been the poor pathophysiological relevance of the source material. Most studies rely on EV derived from immortalized cell lines or primary cells cultured *in vitro*, often exposed to a single artificial stimulus (13, 14). This reductionist approach fails to recapitulate the complexity of the tissue microenvironment and the diverse signals present during pathological processes such as myocardial infarction. In contrast, our approach isolates primary CM-EVs generated *in vivo* in response to the full spectrum of stimuli occurring during cardiac injury, including hypoxia, apoptosis, inflammation, and mechanical stress.

Proteomic analysis of isolated CM-EVs confirmed their cardiomyocyte origin, revealing enrichment in well-characterized myocardial proteins such as myosin heavy chain 6 (Myh6) and other sarcomeric components, consistent with prior evidence that cardiomyocyte injury releases troponin-containing EV into the circulation (15). Importantly, our approach enables systematic molecular characterization of CM-EVs cargo across multiple levels (proteomics, transcriptomics, lipidomics), allowing researchers to construct a comprehensive “ID card” for any EV population of interest. This offers a unique opportunity to interrogate how disease-modifying conditions alter EV composition and, in turn, their biological activities and downstream targets.

Our method is also highly adaptable for functional studies. Building on previous observations (16), we demonstrated that CM-EVs can transfer biologically active Cre recombinase to target cells: exposure of mTmG macrophages to purified CM-EVs resulted in recombination and GFP expression, confirming delivery of active Cre into recipient cell nuclei.

For this demonstration, it is worth noting that target cells (peritoneal macrophages) were incubated with CM-EVs at a very low ratio (100:1) for 24 h, then gently washed and cultured in conventional growth media for another 72h. This setting was designed to ensure minimal residual GFP from CM-EVs remained in the wells or within macrophages. Indeed, EV-mediated GFP transfer to target cells was proven to be very low yield after 24h (17, 18), even under experimental conditions using substantially higher EV doses than those employed in the present study (18). Taking into account the relative short half-life of transferred GFP and our very low quantity of CM-EVs incubated with target cells, we reasoned that any contaminating, CM-EVs-derived GFP detected after 96h would be negligible. Our demonstration therefore provides another direct evidence that primary CM-EVs mediate functional cargo transfer (4). Notably, this effect was observed at relatively low EV-to-cell ratios, underscoring the potency of these primary CM-EVs. Whether this efficiency is unique to CM-EVs or a broader feature of primary EV warrants further comparative study.

Beyond *in vitro* applications, our method opens the door to *in vivo* functional studies of EV derived from a single cell type. Although not shown here, purified CM-EVs can be used to test hypotheses regarding their biodistribution (19) and roles in intercellular communication in specific pathophysiological settings such as myocardial infarction. Prior studies have proposed that cardiac EV modulate neuroinflammation via miR-21(20) or influence systemic immune responses (6, 21), but have lacked the tools to test these mechanisms directly using purified, cell-specific vesicles. Crucially, our approach is not limited to cardiomyocytes: by leveraging the versatility of Cre-driver mouse lines, it can be generalized to isolate EV from virtually any cell type in a tissue-specific and temporally controlled manner. This opens new avenues for dissecting the cellular origin and function of EV in contexts ranging from neurodegeneration or cancer to inflammation and tissue repair.

Several methodological considerations warrant discussion. First, reliance on transgenic Cre-reporter mouse models may restrict immediate application in non-transgenic species or certain non-rodent model systems. Second, the fidelity of Cre-driven GFP labeling inherently depends on the specificity and potential leakiness of the chosen promoter. We mitigated this concern by employing the well-validated, cardiac-restricted Myh6-Cre driver line, confirming exclusive GFP expression in cardiomyocytes via multiple validation methods. Additionally, while this study primarily focused on large EVs (lEVs), the underlying genetic-labeling framework is directly adaptable to small EVs (sEVs) through minor isolation protocol adjustments. Notably, by defining cellular origin genetically, this platform bypasses the known pitfalls of direct surface antibody enrichment, such as off-target binding, cross-reactivity, and variable epitope accessibility (22–24). Our method satisfies the criteria for EV identity, purity, and source attribution, and provides a template for the rigorous characterization urged by the EV community (12, 25).

In summary, we present a versatile, scalable, and genetically tractable method for the selective isolation and functional study of primary, cell-type-specific EV from diseased tissues. This approach enables, for the first time, direct interrogation of the pathophysiological role of a homogeneous EV population in a complex *in vivo* setting. By overcoming key technical barriers in EV research, it should facilitate mechanistic insight, enhance translational relevance, and unlock new therapeutic opportunities. We hope that this strategy will be broadly adopted to dissect EV biology with unprecedented specificity and resolution.

## Author contributions

L.B.: methodology, investigation, analysis, writing original draft. A.R.: methodology, investigation, analysis. P.A.: investigation. V.D.: investigation. M.A.: investigation. L.P.: investigation. L.F.: investigation. C.K.: methodology. E.R.: methodology. F.D.: methodology. D.L.: methodology. X.L.: funding acquisition. J.S.S.: manuscript review & funding acquisition. C.M.B.: funding acquisition. S.M.C.: conceptualization, methodology, analysis, writing original draft, review, supervision & funding acquisition.

## Data availability statement

Data supporting the findings of this study are available from the corresponding author upon reasonable request.

## References

1. Buzas EI. The roles of extracellular vesicles in the immune system. Nat Rev Immunol. 2023;23(4):236–50.

2. Kalluri R, McAndrews KM. The role of extracellular vesicles in cancer. Cell. 2023;186(8):1610–26.

3. Boulanger CM, Loyer X, Rautou PE, Amabile N. Extracellular vesicles in coronary artery disease. Nat Rev Cardiol. 2017;14(5):259–72.

4. Mathieu M, Martin-Jaular L, Lavieu G, Thery C. Specificities of secretion and uptake of exosomes and other extracellular vesicles for cell-to-cell communication. Nat Cell Biol. 2019;21(1):9–17.

5. Loyer X, Zlatanova I, Devue C, Yin M, Howangyin KY, Klaihmon P, et al. Intra-Cardiac Release of Extracellular Vesicles Shapes Inflammation Following Myocardial Infarction. Circ Res. 2018;123(1):100–6.

6. Hegyesi H, Pallinger E, Mecsei S, Hornyak B, Kovacshazi C, Brenner GB, et al. Circulating cardiomyocyte-derived extracellular vesicles reflect cardiac injury during systemic inflammatory response syndrome in mice. Cell Mol Life Sci. 2022;79(2):84.

7. Martins-Marques T, Girao H. The good, the bad and the ugly: the impact of extracellular vesicles on the cardiovascular system. J Physiol. 2023;601(22):4837–52.

8. Kowal J, Arras G, Colombo M, Jouve M, Morath JP, Primdal-Bengtson B, et al. Proteomic comparison defines novel markers to characterize heterogeneous populations of extracellular vesicle subtypes. Proc Natl Acad Sci U S A. 2016;113(8):E968–77.

9. Mathieu M, Nevo N, Jouve M, Valenzuela JI, Maurin M, Verweij FJ, et al. Specificities of exosome versus small ectosome secretion revealed by live intracellular tracking of CD63 and CD9. Nat Commun. 2021;12(1):4389.

10. de Voogt WS, Frunt R, Leandro RM, Triesscheijn CS, Monica B, Paspali I, et al. EV-Elute: A universal platform for the enrichment of functional surface marker-defined extracellular vesicle subpopulations. J Extracell Vesicles. 2024;13(12):e70017.

11. Ackers-Johnson M, Li PY, Holmes AP, O’Brien SM, Pavlovic D, Foo RS. A Simplified, Langendorff-Free Method for Concomitant Isolation of Viable Cardiac Myocytes and Nonmyocytes From the Adult Mouse Heart. Circ Res. 2016;119(8):909–20.

12. Welsh JA, Goberdhan DCI, O’Driscoll L, Buzas EI, Blenkiron C, Bussolati B, et al. Minimal information for studies of extracellular vesicles (MISEV2023): From basic to advanced approaches. J Extracell Vesicles. 2024;13(2):e12404.

13. van Niel G, Carter DRF, Clayton A, Lambert DW, Raposo G, Vader P. Challenges and directions in studying cell-cell communication by extracellular vesicles. Nat Rev Mol Cell Biol. 2022;23(5):369–82.

14. Dixson AC, Dawson TR, Di Vizio D, Weaver AM. Context-specific regulation of extracellular vesicle biogenesis and cargo selection. Nat Rev Mol Cell Biol. 2023;24(7):454–76.

15. Barile L, Moccetti T, Marban E, Vassalli G. Roles of exosomes in cardioprotection. Eur Heart J. 2017;38(18):1372–9.

16. Zomer A, Maynard C, Verweij FJ, Kamermans A, Schafer R, Beerling E, et al. In Vivo imaging reveals extracellular vesicle-mediated phenocopying of metastatic behavior. Cell. 2015;161(5):1046–57.

17. Bonsergent E, Lavieu G. Content release of extracellular vesicles in a cell-free extract. FEBS Lett. 2019;593(15):1983–92.

18. Bonsergent E, Grisard E, Buchrieser J, Schwartz O, Thery C, Lavieu G. Quantitative characterization of extracellular vesicle uptake and content delivery within mammalian cells. Nat Commun. 2021;12(1):1864.

19. Valkov N, Das A, Tucker NR, Li G, Salvador AM, Chaffin MD, et al. SnRNA sequencing defines signaling by RBC-derived extracellular vesicles in the murine heart. Life Sci Alliance. 2021;4(12).

20. Li Q, Hamdalla RH, Dhyani N, Sun L, Gao L, Rudebush TL, et al. Cardiac Injury Regulates Neuroinflammation Through Extracellular Vesicle-Mediated Heart-Brain Crosstalk. JACC Basic Transl Sci. 2025;10(7):101307.

21. Hoyer FF, Naxerova K, Schloss MJ, Hulsmans M, Nair AV, Dutta P, et al. Tissue-Specific Macrophage Responses to Remote Injury Impact the Outcome of Subsequent Local Immune Challenge. Immunity. 2019;51(5):899–914 e7.

22. Kirkpatrick P. Specificity concerns with antibodies for receptor mapping. Nat Rev Drug Discov. 2009;8(4):278.

23. Baker M. Reproducibility crisis: Blame it on the antibodies. Nature. 2015;521(7552):274-6.

24. Ayoubi R, Ryan J, Biddle MS, Alshafie W, Fotouhi M, Bolivar SG, et al. Scaling of an antibody validation procedure enables quantification of antibody performance in major research applications. Elife. 2023;12.

25. Welsh JA, Arkesteijn GJA, Bremer M, Cimorelli M, Dignat-George F, Giebel B, et al. A compendium of single extracellular vesicle flow cytometry. J Extracell Vesicles. 2023;12(2):e12299.

